# Restoring ventilatory control using an adaptive bioelectronic system

**DOI:** 10.1101/488577

**Authors:** Ricardo Siu, James J. Abbas, Brian K. Hillen, Jefferson Gomes, Stefany Coxe, Jonathan Castelli, Sylvie Renaud, Ranu Jung

## Abstract

Electrical stimulation of the diaphragm muscle or phrenic nerve, or ventilatory pacing, serves as an alternative to mechanical ventilation.
Currently available ventilatory pacing systems are open-loop, requiring long set-up sessions and frequent tuning.
A neuromorphic closed-loop bioelectronic controller capable of autonomously adapting current amplitude to achieve a desired breath volume profile has been developed.
The controller was able to achieve a desired volume profile in intact animals and restore tidal volume to values observed prior to injury in spinal cord hemisected animals.
This controller architecture allows for ventilatory pacing that requires minimal technician input during setup and constantly adapts to account for changes such as those induced by muscle fatigue.
The adaptive control system could be used as a respiratory rehabilitation tool to strengthen inspiratory muscles and for automated weaning from mechanical ventilation.

**Abstract:** Ventilatory pacing via electrical stimulation of the phrenic nerve or of the diaphragm has been shown to enhance quality of life compared to mechanical ventilation. However, commercially-available ventilatory pacing devices require initial manual specification of stimulation parameters and frequent adjustment to achieve and maintain suitable ventilation over long periods of time. Here, we have developed an adaptive, closed-loop, neuromorphic, pattern-shaping controller capable of automatically determining a suitable stimulation pattern and adapting it to maintain a desired breath volume profile on a breath-by-breath basis. *In vivo* studies in anesthetized intact and C2-hemisected male Sprague-Dawley rats indicated that the controller was capable of automatically adapting stimulation parameters to attain a desired volume profile. Despite diaphragm hemiparesis, the controller was able to achieve a desired volume in the injured animals that did not differ from the tidal volume observed prior to injury (*p*=0.39). The closed-loop controller was developed and parametrized in a computational testbed prior to *in-vivo* assessment. This bioelectronic technology could serve as an individualized and autonomous respiratory pacing approach for support or recovery from ventilatory deficiency.

## Introduction

When the autonomic control of ventilation is compromised, such as with cervical spinal cord injury or central hypoventilation syndrome, means of artificial ventilation are often needed. Dependence on mechanical ventilation to treat hypoventilation can cause a considerable impact on quality of life. Although mechanical ventilators are widely used to treat respiratory insufficiency (Kacmarek, 2011), mechanical ventilation can in itself have detrimental effects on lung health (Claxton *et al*., 1998; Ricard *et al*., 2003). Positive pressure mechanical ventilators have been reported to cause alveolar damage and lead to diaphragm muscle atrophy, with studies indicating onset of atrophy after 18 hours of mechanical ventilation (Andrew Shanely *et al*., 2002; Ricard *et al*., 2003; Levine *et al*., 2008; Zambon *et al*., 2016).

Restoration of ventilation through electrical stimulation (pacing) is an alternative to mechanical ventilators in cases where the phrenic nerve is intact, but the pre-motor neurons have been damaged or when the respiratory central pattern generator (CPG) has been compromised. Pacing can be accomplished through stimulating cuff electrodes on the phrenic nerve, intramuscular electrodes in the diaphragm muscle, or catheter-based transvenous electrodes (Reynolds *et al*., 2017). A series of stimulating pulses delivered to the target cause contraction of the diaphragm muscle and thus elicit a functional breath (Glenn *et al*., 1988; Chervin & Guilleminault, 1994; DiMarco, 1999, 2009; Onders, 2012). Pacing approaches have the benefit of neural activation of the diaphragm to produce negative pressure ventilation, eliciting a breath in a more physiological manner. Pacing could also slow or reverse the development of muscle atrophy, and possibly promote central neuroplasticity for improved recovery (Onders *et al*., 2007; DiMarco, 2009; Onders, 2012; Masmoudi *et al*., 2017).

Current commercially-available systems operate in an open-loop manner, in which the clinician must determine and set stimulation parameters that are suitable for ventilatory pacing for a given individual (Onders, 2012). However, there is a need to adjust the stimulation parameters for pacing at a short time-scale in response to physiological and metabolic demand, postural load changes (Winslow & Rozovsky, 2003), and stimulation-induced muscle fatigue. Changes over the long time-scale such as electrode encapsulation or muscle atrophy, should also be accounted for.

The limitations of an open-loop system could be addressed by an adaptive closed-loop bioelectronic control system that adjusts stimulation to provide sufficient ventilation to the user. To achieve efficient and responsive ventilation, the controller should be able to automatically and repeatably produce a suitable breath volume profile and adapt respiratory rate with minimal technician intervention. As an initial step to develop an adaptive ventilatory controller for diaphragmatic pacing, the aims of the present study were the design and development of a neuromorphic closed-loop adaptive controller capable of achieving and maintaining a desired breath volume profile at a pre-determined fixed respiratory rate.

An adaptive ventilatory pacing controller, shown in Figure 1, was developed following a neuromorphic pattern generator/pattern shaper (PG/PS) architecture (Abbas & Triolo, 1997; Kim *et al*., 2009; Jung, 2018).A computational model for the chest and diaphragm in the adult rat was used for initial development of the controller and to determine suitable controller parameters for the PS controller. A rodent model of incomplete spinal cord injury was used to assess the ability of the controller to ameliorate hypoventilation. *In vivo* experiments on both intact and cervical (C2) hemisected animals confirmed that PS-controlled adaptive pacing could achieve and maintain a desired volume profile. In hemisected animals, tidal volume was restored to values observed prior to the hemisection. Pacing also caused a decrease in the elevated end-tidal CO_2_ (etCO_2_) values that resulted from the reduced ventilation after hemisection. Because breathing itself is dynamic, the development of a similarly dynamic neuromorphic controller for ventilation represents an important step in expansion of neurotechnology that is able to, ultimately, replicate and replace autonomic function after ventilatory impairment.

**Figure 1.**
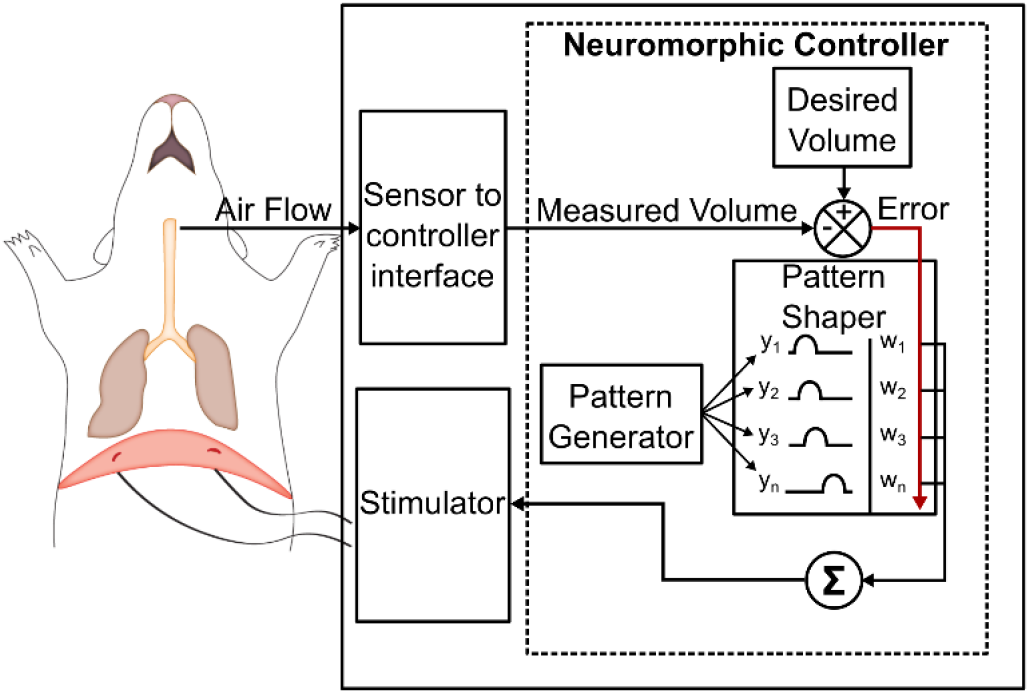
Closed-loop controller architecture for use in experimental studies for diaphragmatic control. The pattern generator sets a fixed respiratory cycle period. The pattern shaper neural network adapts as a function of the instantaneous error between the desired and measured breath volume profiles. The summation of time-shifted neuronal outputs from the pattern shaper dictate controller output to the stimulator.

## Methods

### Animal care and surgical procedure

*In vivo* studies were conducted on spontaneously breathing, adult, male Sprague Dawley rats (open loop stimulation *n = 1*, adaptive stimulation intact *n = 9*, hemisected *n = 7*; 430 ± 60 g) with the approval of the Institutional Animal Care and Use Committee of Florida International University. Rats were initially anesthetized with intraperitoneal pentobarbital (50 mg/kg) followed by supplemental inhaled isoflurane (1.0 – 2.5%) in 100% O_2_. The plane of anesthesia was assessed via a toe-pinch reflex. Body temperature was maintained between 36-38 °C via a heating lamp. A pulse oximeter sensor monitored SpO_2_. A tracheal tube was inserted after a tracheostomy and sutured to the trachea. A small animal pneumotachometer (PTM Type HSE-73-0980, Harvard Apparatus, Holliston, MA) connected directly to the tracheal tube measured air flow. Flow was integrated (PI-1000, CWE Inc., Ardmore, PA) to obtain breath volume. A capnograph (CapStar-100, CWE Inc., Ardmore, PA) monitored end-tidal CO_2_ (etCO_2_) throughout the experiment. 30G stainless steel needle electrodes were inserted subcutaneously to monitor cardiac activity via electrocardiogram. Stainless steel wire intramuscular recording electrodes (SS-304, 44 AWG, AM-Systems, Carlsborg, PA) were inserted below the tongue for recording of genioglossus (GG) muscle activity in some animals (n=2). GG activity was recorded to assess synchrony between the intrinsic inspiratory drive and paced breaths.

A ventral incision provided access to the caudal aspect of the diaphragm muscle. A handheld stimulator (DigiStim 3 plus, Neuro Technologies) with a blunted needle delivered constant current pulses at 1 Hz to map the location of the neuromuscular junction. Stimulation electrodes were implanted near this location. The peritoneal incision remained open to provide visualization of the diaphragm during assessment of muscle twitch thresholds, after which the incision was sutured closed.

A lateral C2 hemisection was performed prior to stimulation in 7 rats to use as an incomplete spinal cord injury (iSCI) model. A dorsal C2 laminectomy was performed to expose the cervical spinal cord. Once exposed, a microscalpel was used to perforate the dura and to sever the left half of the spinal cord at the C2 level. Observing a considerable decrease in tidal volume provided a functional assessment of a proper hemisection. Spinal tissue was also collected for histological assessment to ensure that the hemisection resulted in severance of all descending phrenic pathways.

### Diaphragmatic stimulation

Diaphragmatic pacing was conducted using single stranded, stainless steel electrodes (SS-304, 44 AWG, AM-Systems, Carlsborg, PA) implanted through the abdominal fascia of the diaphragm or through the muscle itself with the use of a suture needle (10 mm, 3/8th circle, cutting, Havel’s Inc., Cincinnati, OH). Stimulation was performed using a constant current programmable stimulator (FNS-16, CWE Inc., Ardmore, PA) delivering cathodic first, biphasic current pulses of 200 μs/phase at 75Hz with amplitude modulated every 40 ms by the adaptive PS controller. The 75 Hz stimulation frequency was shown to provide tetanic contractions in rat skeletal muscle (Jung *et al*., 2009; Fairchild *et al*., 2010). Pulses delivered at 75 Hz were also found to consistently elicit effective tetanic contractions of the diaphragm during preliminary trials and thus this pulse frequency was maintained for all trials in this study. In all experimental trials the initial current pulse amplitude was set to zero which automatically changed once the PS controller was initiated. A maximum limit for current amplitude was set at 4 times the twitch threshold amplitude during open-loop stimulation. Maximum current amplitude values for the stimulated hemi-diaphragms ranged from 0.5 mA to 4.0 mA; the FNS-16 stimulator resolution for the current amplitude is 0.001 mA.

### Experimental protocol

To assess the controller’s ability to achieve a desired ventilatory profile in both intact and hemisected rats, the controller was set to modulate stimulation to match a desired breath volume profile with a constant cycle period. The desired breath volume profile and cycle period were obtained from baseline ventilatory data recorded for at least one minute prior to stimulation in each trial in intact animals and in multiple intermittent recordings prior to injury for the trials in hemisected animal. The desired breath volume profile was defined by averaging all recorded breathing cycles except for non-breathing behaviors, i.e. sighs.

Since the intrinsic biological respiratory drive was either fully active (in intact animals) or partially active (in hemisected animals), it was necessary to achieve entrainment and synchrony between the intrinsic and paced respiratory cycles. Entrainment is defined as a matching of the intrinsic respiratory frequency to the paced respiratory frequency while synchrony is defined as 1:1 entrainment between the intrinsic and paced respiratory cycles with zero-phase shift in the onset of the breath cycle. In intact animals, this was achieved by setting the volume of the desired breath volume profile to 120% of the baseline volume, thus ensuring a strong paced signal that would entrain the intrinsic pattern. In hemisected animals, the desired breath volume profile was maintained at pre-hemisection values unless an increase in desired breath volume was required to promote in-phase synchronization. The duration of the desired breath cycle was set to match the duration of the breath cycle prior to the trial; if required to improve the likelihood of synchronization, this value was adjusted (within ± 10%). The desired breath volume profile was maintained constant throughout the trial.

In trials in intact animals, each trial consisted of at least one minute of spontaneous breathing, followed by at least 100 cycles of pacing with an increased desired breath volume of 120% of the baseline breath volume. There were one to eight trials per intact animal. In animals with iSCI, prior to hemisection, trials were defined as multiple recordings of one minute each of spontaneous breathing (*n=3* per rat, with the exception of rat 11 where *n=2*). After hemisection, trials were defined as at least one minute of spontaneous breathing followed by at least 200 pacing cycles. There were two to six trials post-hemisection per animal.

Expiratory muscles were not stimulated and therefore expiration was passive. A rest period of 15 minutes was allowed between trials to allow the diaphragm to recover from stimulation-induced fatigue. In hemisected animals, a mechanical ventilator was used to maintain proper ventilation during these rest periods and to maintain etCO_2_ at nominal levels.

### Performance measures

The adaptive controller adapts the stimulation output to the magnitude of the error between the desired volume and the measured volume at a given instant. To quantify the deviation from the desired volume and the achieved volume traces over the portion of the respiratory cycle directly affected by inspiratory muscle activation, the inspiratory root mean square error (iRMSE) % for each pacing cycle was defined as

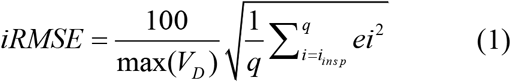

Where *max*(*V_D_*) is the peak value for the desired volume, *i_INSP_* denotes the time step at the start of inspiration, *q* the number of time steps in the inspiratory phase of the desired breath volume profile, and *e_i_* the instantaneous error at time step *i*.

The basis of this error value across several pacing cycles has been used as an indicator of controller accuracy in previous studies regarding evaluation of the PG/PS controller (Abbas & Triolo, 1997; Riess & Abbas, 2000; Stites & Abbas, 2000). In the current study, the number of pacing cycles required for iRMSE % to fall below a certain threshold and maintain it for at least 10 pacing cycles served as an indicator of controller speed of adaptation. A threshold of 5% iRMSE was selected for simulation studies. In animal studies, due to inherent breath-to-breath variability in the inspiratory volume profile, the iRMSE threshold was elevated to 10%. The mean and standard deviation of the iRMSE of the last 40 pacing cycles of the pacing period were used to assess controller performance.

In addition to the iRMSE measures, the effectiveness of pacing in hemisected animals was assessed by evaluating tidal volume and etCO_2_ during pacing. Values for these measures were tabulated under pre-hemisection, post-hemisection without pacing, and post-hemisection with pacing conditions from 40 pacing cycles for tidal volume and 10 pacing cycles for etCO_2_ per condition. The breath cycles post-hemisection without pacing were obtained just prior to initializing pacing, and those post-hemisection with pacing were obtained prior to the end of the 200-pacing cycle pacing period.

### Histology

After completion of the experimental protocols each hemisected animal was perfused with phosphate buffer solution followed by 4% paraformaldehyde. The spinal cord was extracted from the atlas to the C7 vertebra and placed in 4% paraformaldehyde and then transferred to 30% sucrose for dehydration. Spinal cords were then frozen and cryosectioned to obtain 10 μm longitudinal sections. The presence of a gap in the ventrolateral tissue observed under light microscopy was used to indicate a successful hemisection of the ipsilateral descending phrenic pathways.

### Experimental design and statistical analysis

A post-hoc t-test power analysis was performed to assess whether the number of intact (n = 9) and spinal cord hemisected (n = 7) animals provided sufficient power for statistical analysis of the number of pacing cycles required to achieve 10% iRMSE or less. Based on descriptive statistics obtained for this measure (intact = 16.06 ± 6.66 pacing cycles, hemisected = 58.8 ± 27.25 pacing cycles) and an alpha of 0.05, the test provided a power of 0.98 (Df = 14). A repeated measures ANOVA power analysis was also performed post-hoc to assess if 40 pacing cycles was sufficient to identify a difference in tidal volume amongst the pre-hemisection, post-hemisection without pacing, and post-hemisection with pacing conditions in the hemisected rats. A similar analysis was done to determine if 10 pacing cycles of etCO_2_ were sufficient. The resulting power was 1.00 and 0.95 for tidal volume and etCO_2_, respectively. All power analysis was performed using G*Power 3.1.9.2 (Faul *et al*., 2007). Descriptive statistics were obtained using SPSS (IBM, Armonk, NY).

To assess the effect of pacing on tidal volume and etCO_2_ after hemisection, a general linear mixed model (Cnaan *et al*., 1997) was used with fixed effects of trial number and condition (intact, hemisection without pacing, and hemisection with pacing) and random intercept effects of both trial number and measurement occasion (last 40 breath measurements within each trial for tidal volume and last 10 breath measurements for etCO_2_). Both pre-hemisection and post-hemisection with pacing conditions had significantly higher tidal volumes than the post-hemisection without pacing condition (pre-hX vs. post-hX difference = 0.64 ml, p<.0001; post-hXp vs. post-hX difference = 0.59 ml, p<.0001). Pre-hemisection and post-hemisection with pacing conditions did not have significantly different breath volumes (pre-hX vs post-hXp difference = 0.05 ml, p = 0.39). All three conditions were mutually significantly different in etCO_2_ values, with pre-hemisection values lowest, post-hemisection with pacing values higher, and post-hemisection without pacing values higher still (pre-hX vs post-hXp difference = 0.73%, p=.0002; pre-hX vs. post-hX difference = 1.46%, p<.0001; post-hX vs post-hXp difference = 0.73%, p<.0001). The generalized linear mixed model was performed using SAS 9.4 (SAS Institute Inc., Cary, NC).

### PG/PS controller design

The PG/PS controller utilized in this study is modeled on the original PG/PS controller that was designed and used to control cyclic limb movements using functional neuromuscular stimulation. The PG/PS controller had the ability to compensate for muscle fatigue by adapting stimulation pulse parameters and charge delivery to achieve the desired functional outcome (Abbas & Chizeck, 1991; Abbas & Triolo, 1997; Riess & Abbas, 2000; Stites & Abbas, 2000; Ichihara *et al*., 2009; Jung *et al*., 2009; Fairchild *et al*., 2010). The PG module generates the pacing cycle period while the PS module adapts to define stimulation parameters that elicit a desired action. In the current study, a fixed frequency oscillator was used as the PG to produce a fixed respiratory frequency that did not change from breath-to-breath. The PS module consisted of a single-layer neural network with the output profile of each neuron being a raised cosine function; the outputs of the neurons in the set are time-shifted with respect to each other. In its current implementation, the number of neurons was set as 25, with one reaching its peak every 40 ms (which matched the frequency at which the controller sends an update to the stimulator). To create a stimulation pulse train with frequency of 75 Hz, each update to the stimulator resulted in the production of three pulses of identical amplitude. Each neuron reaches its peak activity at a specific time-step but its temporal profile overlaps across a specified number of neighboring neurons. The width of the temporal profile determines the number of overlapping activation profiles and is defined in the controller by the parameter *n_a_*. The output, *z(t)*, of the PS at each time step is the weighted summation of the non-zero neuron outputs at that time step and is given by

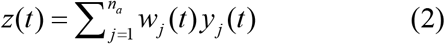

Where the output of a neuron *j* at time *t* is defined as *y_j_*(*t*), with the weight for that specific neuron set as *w_j_*(*t*). The output, *z*(*t*), is restricted to the range of values between 0 and 1 and then scaled by the maximum current amplitude defined for each channel.

Each scaling factor, or weight, for each neuronal output is modified based on an instantaneous error between a pre-established target breath volume cycle profile and the measured volume signal. The controller works in a feedforward manner by modifying each neuron’s weight at each time step based on the measured value of error at that time. The change in weights for neuron *j* is

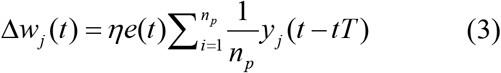

Where *η* is the learning rate, *e*(*t*) is the instantaneous error at time *t, T* is the update period, *n_p_* is number of past activations over which the error is averaged, and *i* the *n_p_* index.

Computer simulations were carried out to elucidate the range of parameters over which adaptation would be stable and errors minimized. These ranges were then used in animal studies to assess *in vivo* implementation of the controller.

### Musculoskeletal diaphragm model

A computational model of diaphragm and chest biomechanics was used as a testbed to design and develop the PG/PS adaptive controller prior to *in-vivo* evaluation. The musculoskeletal model and PG/PS controller were implemented in the LABVIEW programming environment using the Control Design and Simulation module (National Instruments Inc., Austin, TX). The chest model is a biomechanical model in which the mechanical action of diaphragm muscle displacement is assumed to correspond linearly to breath volume (Milhorn, 1966; Lessard, 2009). A proportionality constant, *k_p_*, was implemented to calculate breath volume from muscle displacement. The mechanical action of the diaphragm is defined by the inertance of the chest system, *m_L_*, the elastance of the system, *K_L_*, and the damping coefficient of the system, *B_L_*. Chest biomechanics was described mathematically as a function of diaphragm contractile force via the following equation,

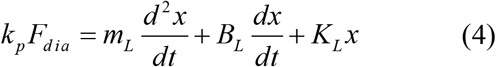

Where the input to the model is the force exerted by the diaphragm muscle, *F_dia_*, and the output is the displacement of the diaphragm, *x*. The mechanical circuit is shown in Figure 2(A).

**Figure 2.**
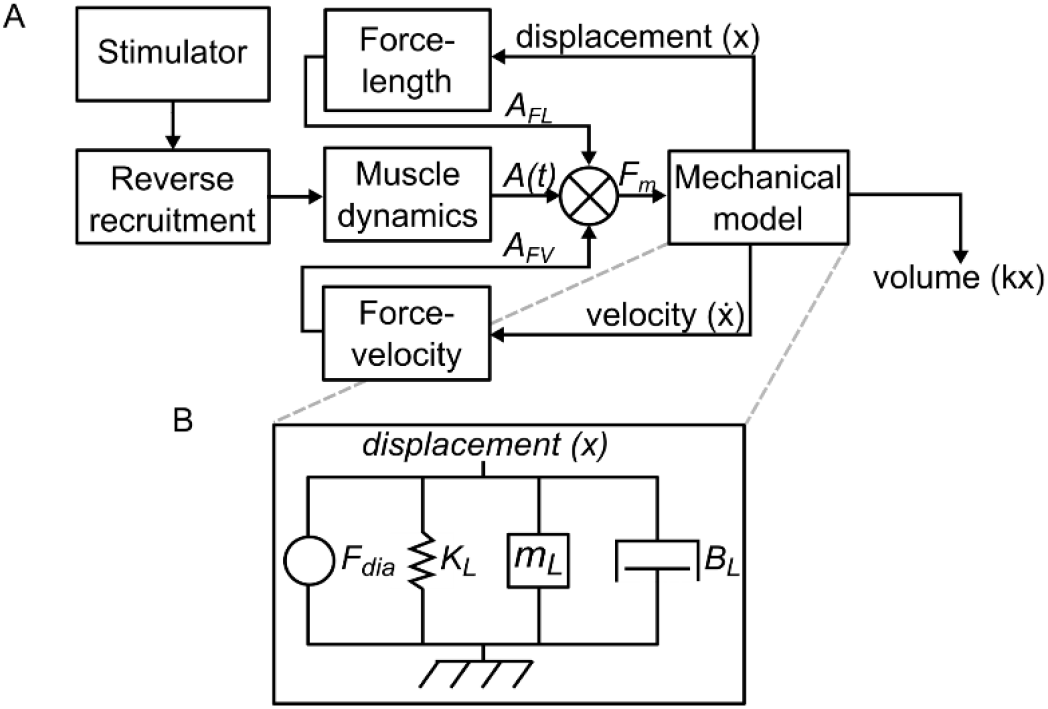
A diaphragm muscle model (A) and chest biomechanical model (B) are used for simulation of closed-loop pacing of the diaphragm muscle. The mechanical model output is the displacement of the diaphragm (*x*), which then translates to volume through a conversion factor (*k*). The displacement and velocity are delivered back into the muscle model to factor into the force-length and force velocity contributions. The controller modulates stimulation delivered based on the error between a desired volume and the output of the model.

The diaphragm muscle model was based off the multiplicative quadriceps muscle model with reverse recruitment shown in Figure 2(B) (Teixeira *et al*., 1991; Abbas & Chizeck, 1991). Briefly, the total force exerted by the muscle, *F_dia_*, is given by the product of a dynamic muscle activation model output *A_k_*, force-length factor *A_FL_*, and a force-velocity factor *A_FV_*.

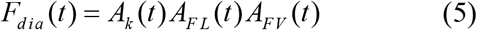

Intrinsically, fatigue-resistant fibers are recruited first while fast-fatigable fibers are recruited later when there is a higher demand for recruitment of muscle fibers. However, due to the nature of extraneural electrical stimulation in which larger fibers have a lower threshold, the larger fast fatigable fibers are recruited before the slow fatigue-resistant fibers are activated resulting in reverse recruitment (Peckham & Knutson, 2005). This reverse recruitment produces nonlinear input/output properties that were modeled using

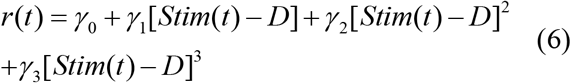

in which muscle recruitment, *r*(*t*), is calculated to mimic the intrinsic recruitment curve observed experimentally in previous studies in rats (Mantilla *et al*., 2010). *D* represents the dead band, or the pulse amplitude below which no fibers are recruited, and the constants, *γ*_0_, *γ*_1_, and *γ*_3_ were modified to obtain a cubic curve that fit these experimental results.

The force generated as fiber recruitment increases is a dynamic process which was modeled using a second-order autoregressive, moving average (ARMA) model with input and output delays (Abbas & Chizeck, 1991).

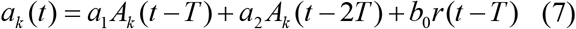

Constants *a*_1_ and *a*_2_ define the linear dynamics of the model, while *b*_0_ defines the muscle input gain. The delay *T*, was based on the model update period of 14 ms.

The force-length component, *A_FL_*, and force-velocity component, *A_FV_*, serve as scaling factors to the force generated by the dynamic model. Force-length factor was estimated with a quadratic equation,

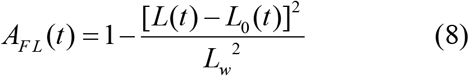

with *L* being the length at instant *t, L_o_* the reference length, and *L_w_* the maximum length of the muscle.

The force-velocity factor was obtained via

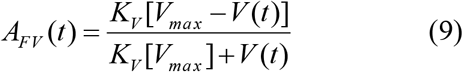

with *K_V_* being the force-velocity slope and *V_max_* the maximum shortening velocity. Constants for all equations and their values can be found in Appendix Table 1.

### Computational trials for parametrization of suitable controller parameters

A previous study has illustrated the ability of the PS controller to adapt across a wide range of controller parameters (Stites & Abbas, 2000). However, in this study the PS was assigned to follow a trajectory for limb movement that differs considerably in profile, amplitude, and duration from one derived from respiratory volume. A LabVIEW implementation of the PS controller and musculoskeletal model was used to carry out the computer simulations. All simulations utilized a predetermined desired trajectory from a baseline volume profile obtained from experimental data.

The controller parameters varied in computer simulation studies were: number of active neurons at any one time-step, number of past neuron activations over which to average the instantaneous error, and learning rate. The controller update frequency was set at 25 Hz and the number of neurons was set to 25. Learning rate values tested were 0.0001, 0.0005, 0.001, 0.002, 0.003, and 0.004. The values tested for the number of active neurons tested ranged from 1 to 19 (in steps of 2) and the values tested for the number of past activations ranged from 2 to 20 (in steps of 2). Each of these simulated trials were done for a length of 100 pacing cycles.

To determine the controller parameters suitable for fast, accurate, and stable performance, only parameter triads that would pass selection criteria were selected. These criteria were: iRMSE below 5% within 20 pacing cycles, mean iRMSE below 5% and a standard deviation below 2.5% iRMSE after <5% iRMSE was attained. Processing of trial results for parameter set selection was performed using MATLAB (Mathworks Inc., Natick, MA).

## Results

### Simulations demonstrate the pattern shaping capabilities of the controller

Using the musculoskeletal model as a developmental testbed, a closed-loop PG/PS controller capable of adapting and modulating the stimulation output to elicit a desired breath volume profile was developed. Results from a sample simulation trial using 3 active neurons with 6 past activation values with a learning rate of 0.001 are presented in Figure 3. In Figure 3(A, B), volume and stimulation traces for the first ten pacing cycles show how the controller can adapt the stimulation levels so that the measured volume profile matches a pre-set desired volume profile. Figure 3(C) shows that in pacing cycles 91-100 the measured and desired volume profiles still match closely. Figures 3(E) and 3(F) show how iRMSE and charge delivered changed over a 100-pacing cycle trial period. When the trial starts, a decrease in iRMSE to a value below 5% iRMSE by pacing cycle # 5 is seen. For the rest of the trial, iRMSE remains well below 5% and finally decreases to an average of 0.96% in the last 10 pacing cycles of the simulation trial. Charge delivered per pacing cycle increases initially as the controller adapts to increase the elicited breath volume profile and match the desired profile. Given that the controller is continuously adapting, charge delivered then decreases as the controller fine tunes the neuronal weights to further reduce iRMSE. Once the error is at a minimum, the charge delivered remains constant for the rest of the trial unless there is a change in the error.

**Figure 3.**
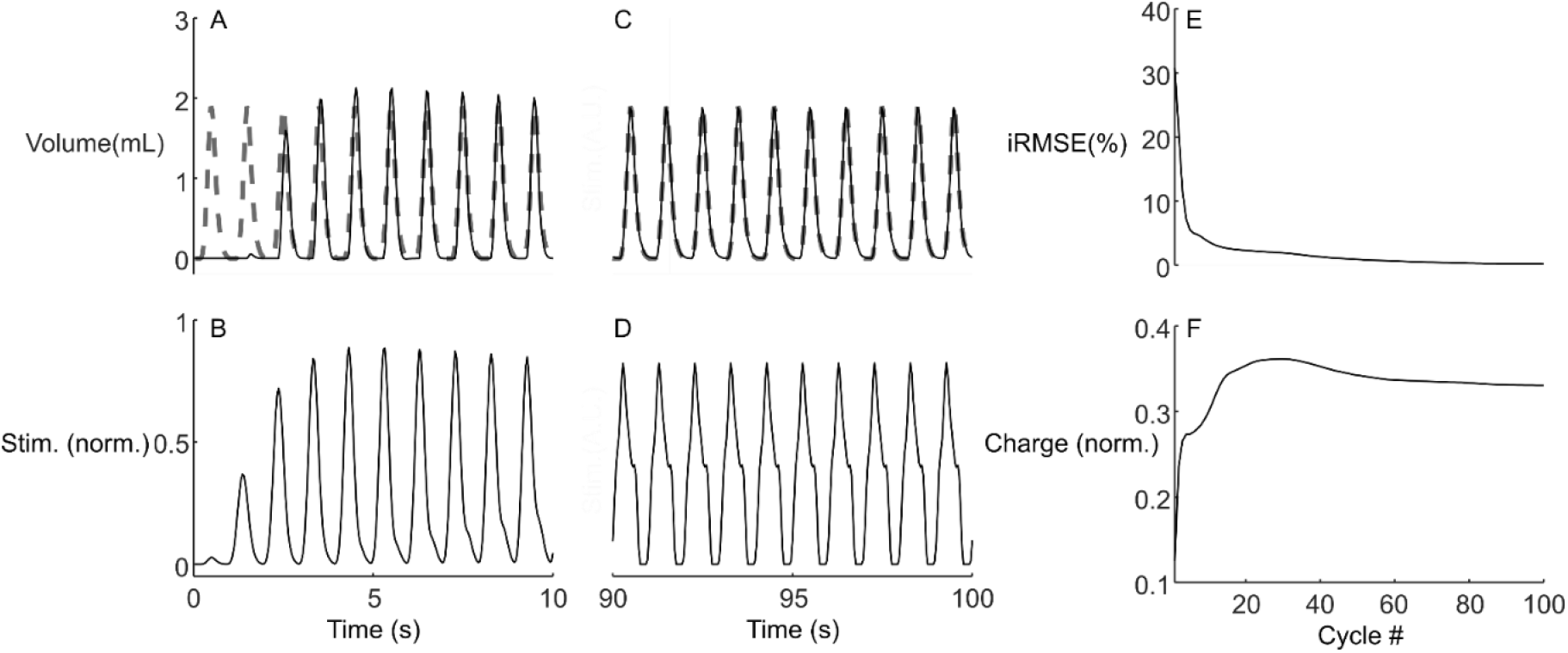
Adaptive PS controller effect in breath volume in a simulated trial. Example of a simulation trial showing the controller adapting the stimulation output (**B, D**) to match the measured volume (**A,C**, solid) to the desired volume (**A,C**, dashed) during the initial stimulation (pacing cycles 1-10) and at end (pacing cycles 91-100) of a 100 second trial.(**E**) Inspiratory RMSE throughout 100 pacing cycles in a simulated trial using controller parameters obtained from parametrization (*η*=0.001, *n_a_* = 3, *n_p_* = 6). (**F**) The corresponding normalized charge delivered by the controller as it adapted the stimulation output to match the targeted volume profile.

### Synchronization between intrinsic breaths and paced breaths *in-vivo*

*In-vivo* trials demonstrated the ability of the controller to adapt to fatigue and reduce inspiratory RMSE in both intact and hemisected anesthetized rats. However, unlike in computational studies, the residual intrinsic respiratory drive interacts with the controller in a manner that significantly affects the controller’s performance. Figure 4, which shows the first 100 seconds of a trial, depicts how the intrinsic respiratory drive interacts with the pacing after it is initiated. Initially stimulation is set to zero as the controller has yet to adapt and increase stimulator output. However, as the error values increase due to the mismatch in timing between the intrinsic breaths and the paced breaths, the controller adapts gradually increasing charge delivered during each pacing cycle. Eventually, the intrinsic breaths synchronize with the pacing induced breaths.

**Figure 4.**
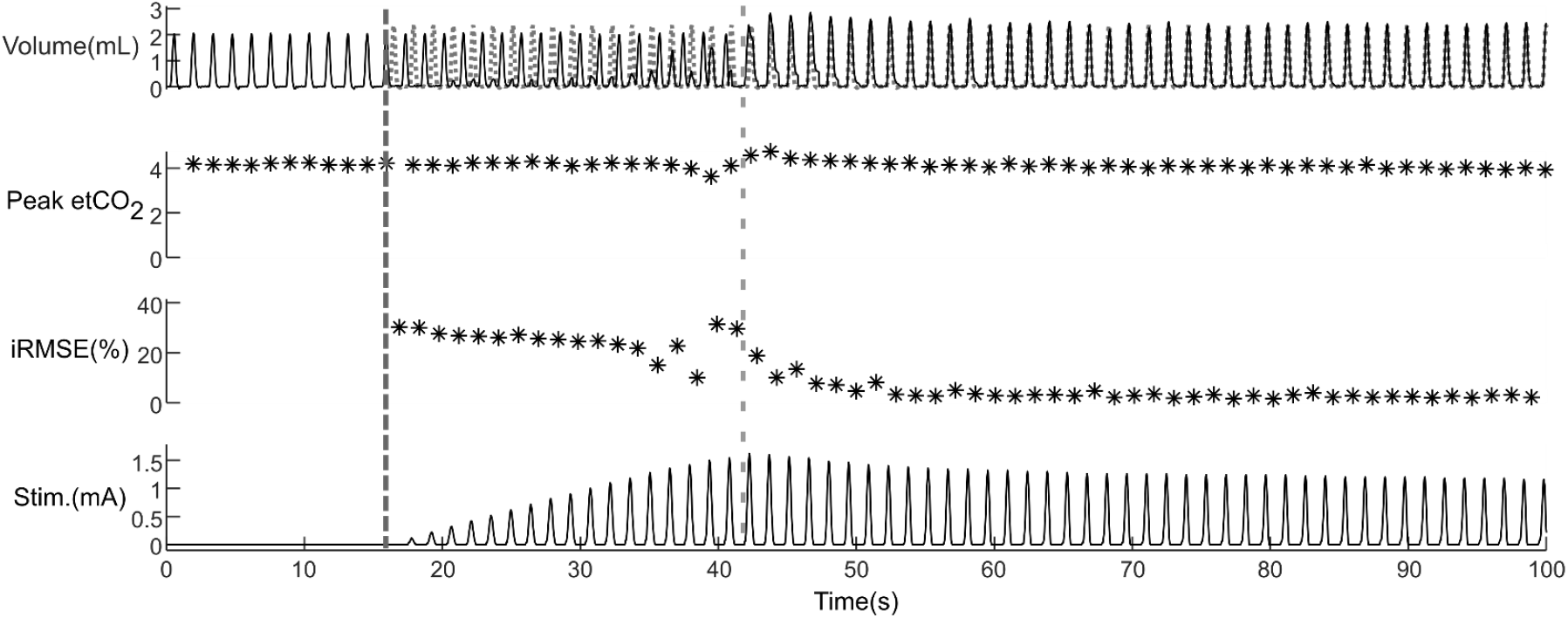
Adaptive PS controller implementation in an intact animal. The controller adapts stimulation output to match the measured volume (solid line) to the desired volume (dotted line). The controller was turned on at 10 seconds (dark grey vertical dashed line) with controller parameters (*n_a_* = 3, *n_p_* = 6, *η* = 0.001). The controller was able to cause successful synchronization after 14 pacing cycles (light grey vertical dashed line) and maintained low iRMSE thereafter.

### Adaptive current modulation is able to achieve and maintain a desired volume profile

After successful adaptation, the measured volume profile closely matches the desired profile, as demonstrated by low iRMSE, and that performance is maintained for the remainder of the trial (Figure 5). Figure 5(A) – 5(B) shows a comparison of the volume profile between a breath elicited using an open loop approach with a square-shaped burst and a breath elicited using pulse amplitudes determined by the PS controller after adaptation. The open-loop approach in Figure 5(A), other than lacking adaptation to extrinsic factors, elicits a breath that differs significantly from a natural breath.

**Figure 5.**
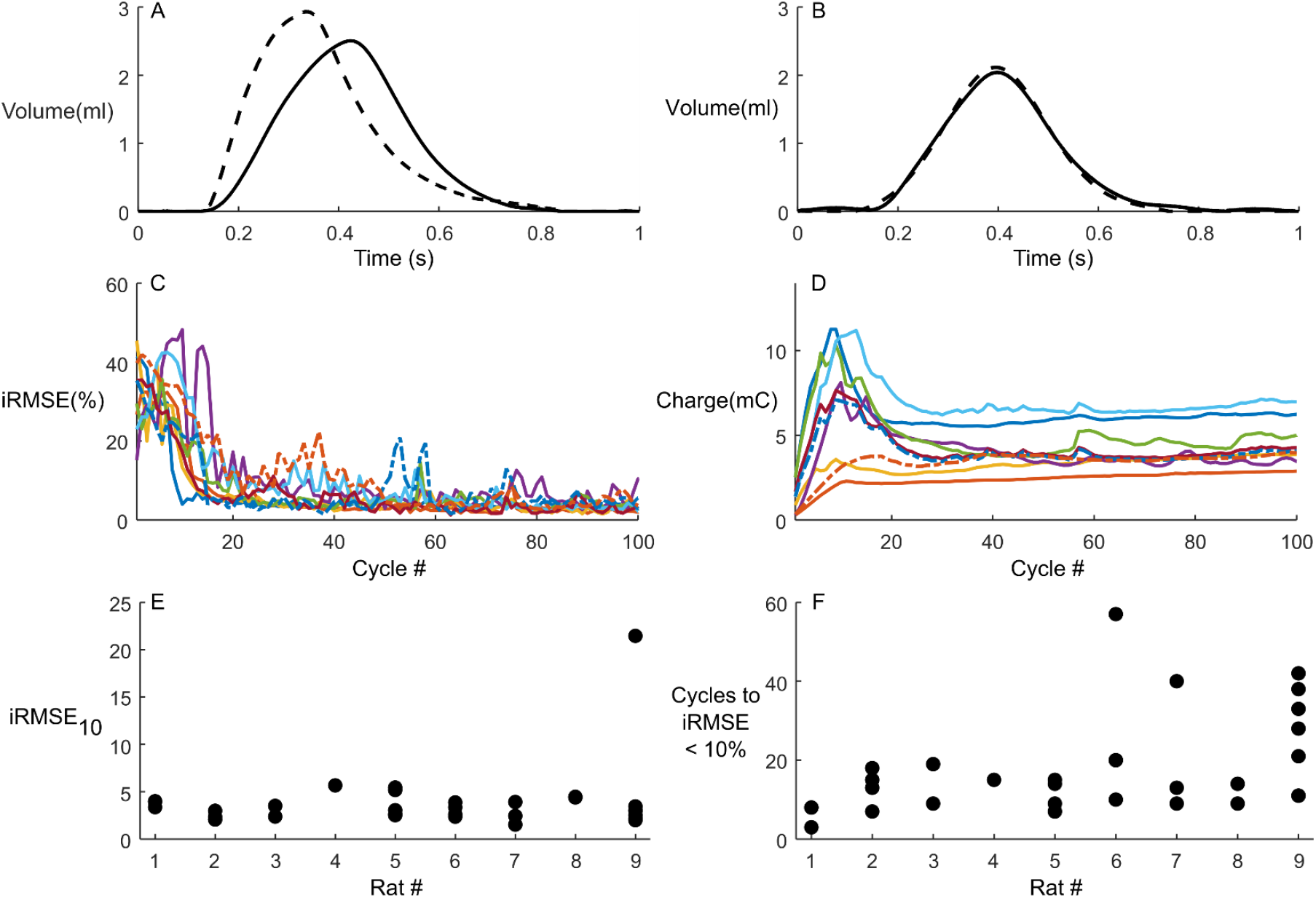
Ability of adaptive PS control to achieve a desired breath volume profile in intact animals. (**A**) Open-loop stimulation using a square-shaped stimulation burst elicited a breath profile (solid line) different from that of a spontaneous breath (dashed line). (**B**) The adaptive controller modulated stimulation to elicit a breath profile similar to a desired breath profile derived from a spontaneous breath. (**C**) Average inspiratory Root Mean Square Error (iRMSE) between the desired and measured breath volume profiles for each animal. iRMSE is elevated at the initiation of the pacing due to lack of synchrony between the breaths produced during intrinsic respiration and those elicited by pacing. After synchrony, the controller is able to reduce and maintain a low iRMSE across all animals. (**D**) Average charge delivered per pacing cycle per animal. The controller tended to trigger entrainment by increasing stimulation output until synchronization occurred. (**E**) Average iRMSE in the last 10 pacing cycles per trial for all animals. iRMSE10 was maintained below 10% except for one trial for animal 9, in which a desynchronization occurred during the last 10 pacing cycles. (**F**) Number of pacing cycles per trial required to achieve the desired 10% threshold, including the pacing cycles required to reach synchronization.

However, the breath elicited by the adaptive PS controller in Figure 5(B) matches the desired breath closely.

A summary of the iRMSE and charge per pacing cycle for all intact animal trials is presented in Figures 5(C) and 5(D), respectively. Entrainment with synchronization usually occurred by pacing cycle 20 and, within 5 pacing cycles, this resulted in an iRMSE of 10%. As seen in Figure 5(C), iRMSE remained below 10% for most of the trials, with the exceptions of periods where intrinsic perturbations occurred. These events are marked by a sudden rise and drop in iRMSE. In seven trials, pacing was performed for an extended duration of up to 15 minutes (not shown). These extended trials also showed iRMSE below 10% throughout the trial when in synchrony. As shown in Figure 5(D), charge delivered per pacing cycle increased substantially until entrainment and synchronization occurred. After this initial period, the controller fine-tuned the stimulator output to obtain the desired volume profile. The charge delivered remained relatively steady for the rest of the trial, changing only when intrinsic perturbations occur.

Figures 5(E) and 5(F) show a summary of the data from Figure 5(C). Figure 5(E) shows the average number of pacing cycles each animal took to achieve the 10% iRMSE threshold and maintain it for at least 10 pacing cycles. This value is heavily dependent on how fast and robustly the intrinsic respiratory rhythm entrains to the pacing.

Given that some of the factors that affect entrainment and synchronization differed between animals, the variability in this value was high. However, the number of pacing cycles required to reach this value among all intact animals were 16.06 ± 6.66 pacing cycles.

Figure 5(F) shows the average iRMSE % for the last ten pacing cycles of the 100-pacing cycle trials for each animal (iRMSE10). On one trial with animal 9, desynchronization occurred once during this ten-pacing cycle period; this resulted in the outlier observed. All other trials remained with consistently low iRMSE10 for the last ten pacing cycles across all trials.

In C2-hemisected animals, the adaptive controller was able to elicit breath volume profiles that were comparable to those observed prior to the injury. Figure 6 shows the initial 100 seconds of a trial showing the adaptive pacing’s effect in a hemisected animal with the target breath volume being the breath volume profile collected pre-hemisection. Despite a considerable reduction in the rodent’s breath volume with only the impaired intrinsic drive, the controller could adapt to sufficiently stimulate the diaphragm muscle to attain the breath volume profile observed pre-hemisection. This breath volume profile was maintained for the duration of the trial. Nine trials across multiple animals were extended to last over 25 minutes to assess longer periods of ventilatory pacing (not shown). Except for periods of desynchronization between the paced and intrinsic breaths, iRMSE remained below 10% in these extended trials.

**Figure 6.**
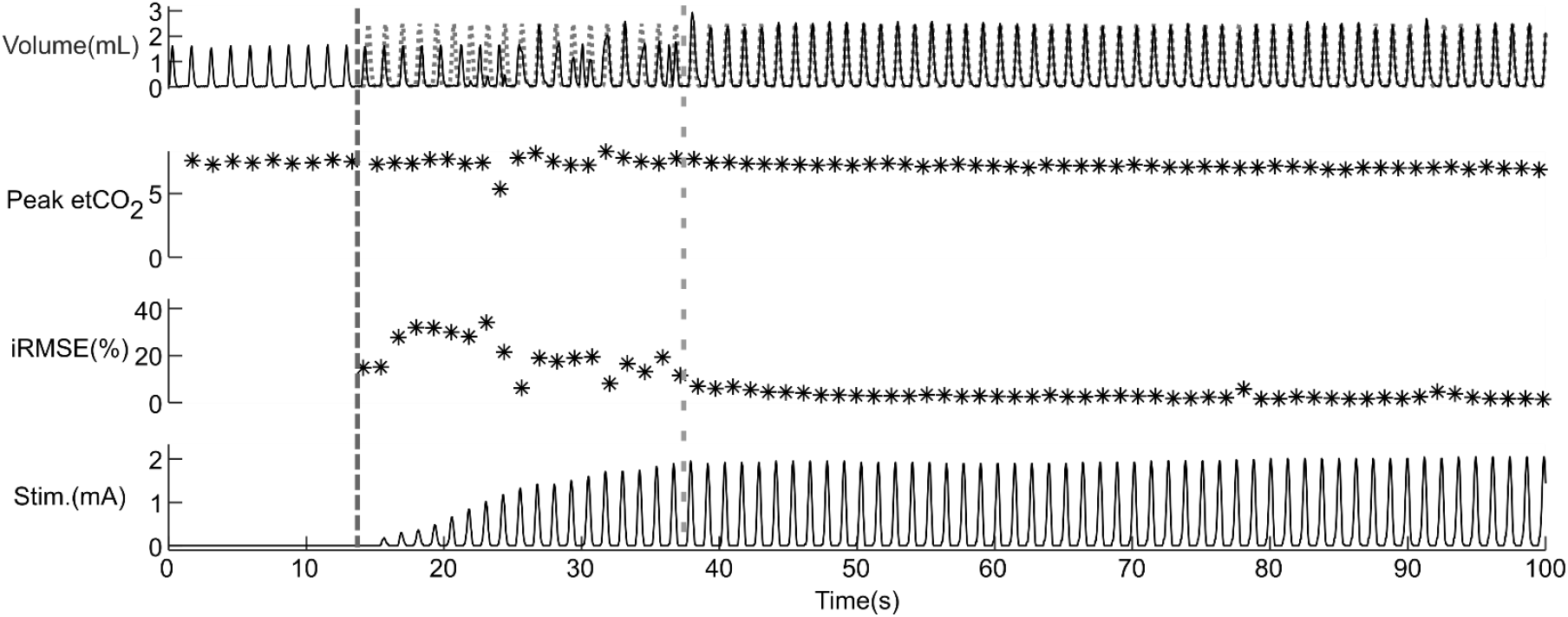
Adaptive PS controller implementation in a hemisected animal. The dark grey dashed line depicts controller initiation. The rodent is spontaneously breathing at a tidal volume of 1.8 ml before stimulation while the desired tidal volume was set to 2.5 ml. After the 20 pacing cycles required for entrainment and synchronization (light grey dashed line), the elicited volume profile quickly matched the desired volume profile (dotted line). This increased volume was maintained for the rest of the rest of the stimulation trial.

Overall controller performance and stimulation results for C2-hemisected animals can be seen in Figure 7 and Figure 8. Across all hemisected animals, iRMSE after entrainment and synchronization remained at 10% or less as seen in Figure 7(A). The number of pacing cycles required to achieve an iRMSE of less than 10% for more than 10 pacing cycles is shown in Figure 7(B). Overall, 58.8 ± 27.25 pacing cycles were required across all hemisected animals to reach 10% iRMSE and maintain it within the 200-pacing cycle trial period. This value exceeded 200 pacing cycles in two trials for animal 16 and thus those trials were excluded from this average. The pacing cycle number to threshold in hemisected animals was higher in comparison to those of intact animals, suggesting that spinal cord injury has a detrimental effect on promotion of respiratory synchronization.

**Figure 7.**
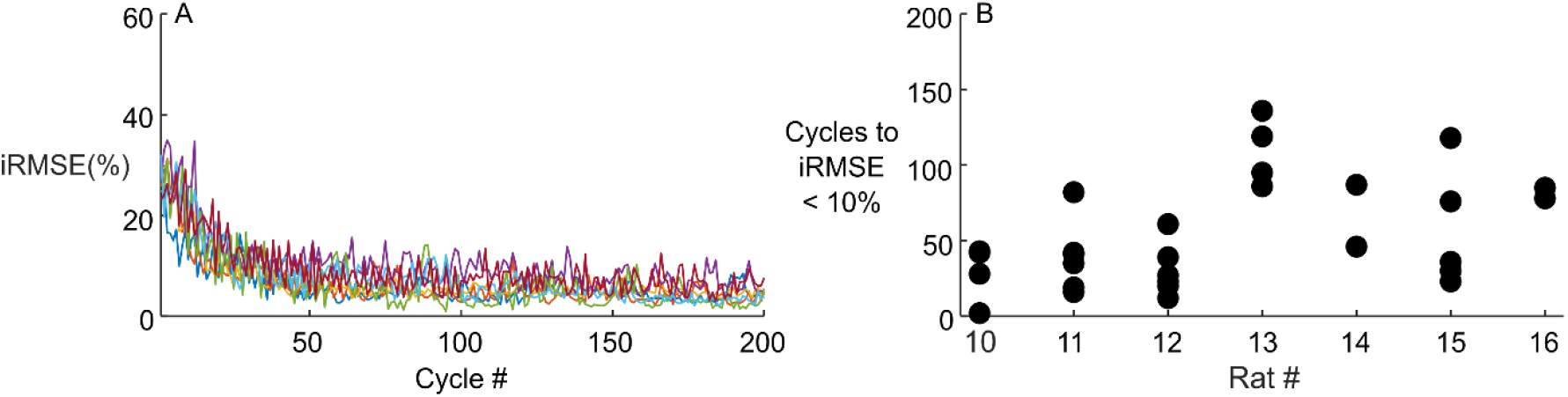
Ability of adaptive PS control to achieve a desired breath volume profile in hemisected animals. Hemisected animals took longer to achieve entrainment and synchrony between the paced and spontaneous breaths than observed in intact animals. This effect can be seen in (**B**) where the iRMSE was not reduced below 10% until very late in the trial in some instances. However, as can be seen in (**A**), iRMSE averaged around 10% for the duration of the trial.

Results across all hemisected animals and across all trials are shown in Figure 8. The effect on tidal volume is shown in Figure 8(A). Tidal volume across all animals was significantly higher during post-hemisection with pacing (post-hXp) than that during post-hemisection without pacing (post-hX) for the last 40 pacing cycles of each trial (0.59 ml increase). Pre-hemisection (pre-hX) tidal volume was not significantly different from post-hemisection with pacing (0.05 ml difference). This shows the controller’s ability to closely match a desired volume profile despite injury.

**Figure 8.**
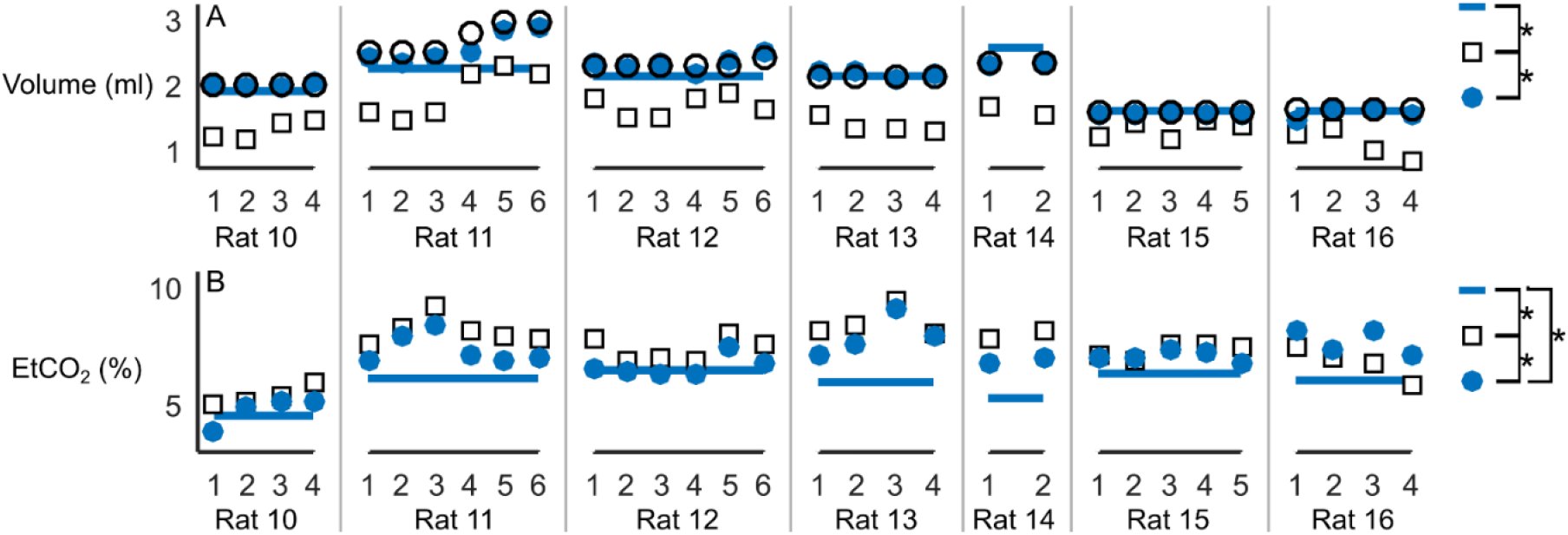
Effect of pacing on tidal volume and end-tidal CO_2_ across trials for all hemisected animals. Tidal volume and etCO_2_ pre-hemisection (pre-hX; solid line), post-hemisection without pacing (post-hX; empty square), and post-hemisection with pacing (post-hXp, solid circle) conditions in each trial for each animal. Pacing was able to restore tidal volume to target values (empty circle) similar or larger than those observed prior to the injury (pre-hX vs. post-hX difference = 0.64 ml, p<.0001; post-hXp vs. post-hX difference = 0.59 ml, p<.0001). EtCO_2_ was also reduced in most cases during post-hemisection with pacing. However, the amount reduced was not sufficiently large to match the etCO_2_ values observed when pre-hemisection values were recorded (pre-hX vs post-hXp difference = 0.73%, p=.0002; pre-hX vs. post-hX difference = 1.46%, p<.0001; post-hX vs post-hXp difference = 0.73%, p<.0001). Pre-hemisection values obtained from n = 3 trials, with the exception of rat 11 (n = 2 trials).

In most trials, pacing had an effect on etCO_2_ as shown in Figure 8(B). All three conditions were mutually significantly different, with pre-hemisection values lowest, post-hemisection with pacing values higher, and post-hemisection without pacing values highest (pre-hX vs post-hX difference = 1.46%; pre-hX vs post-hXp difference = 0.73%; post-hX vs post-hXp difference = 0.73%). While the resulting etCO_2_ was still well above the normal etCO_2_ range, this reduction does show that adaptive pacing is able to have an effect on etCO_2_. It is likely that etCO_2_ can be further reduced to normative values with respiratory rate modulation once etCO_2_ feedback is implemented.

Controller parameters differed to a certain extent amongst intact animals; however, this further shows the robustness of the controller under multiple controller parameters.

Appendix Table 2 shows the controller parameters across all intact animals. Controller parameters for all hemisected animals were set to *n_a_*=3, *n_p_* = 6, and *η* =0.001.

Figure 9(A) shows 15 seconds of a trial in a hemisected rodent in which synchronization was lost. The desired and actual volume profile for each breath are shown in the top row. The muscle activity recorded from the genioglossus (GG) muscle reflects the intrinsic respiratory drive (Brouillette & Thach, 1980; Fregosi & Fuller, 1997; John *et al*., 2005), while the stimulation trace shows the timing at which pacing occurred.

**Figure 9.**
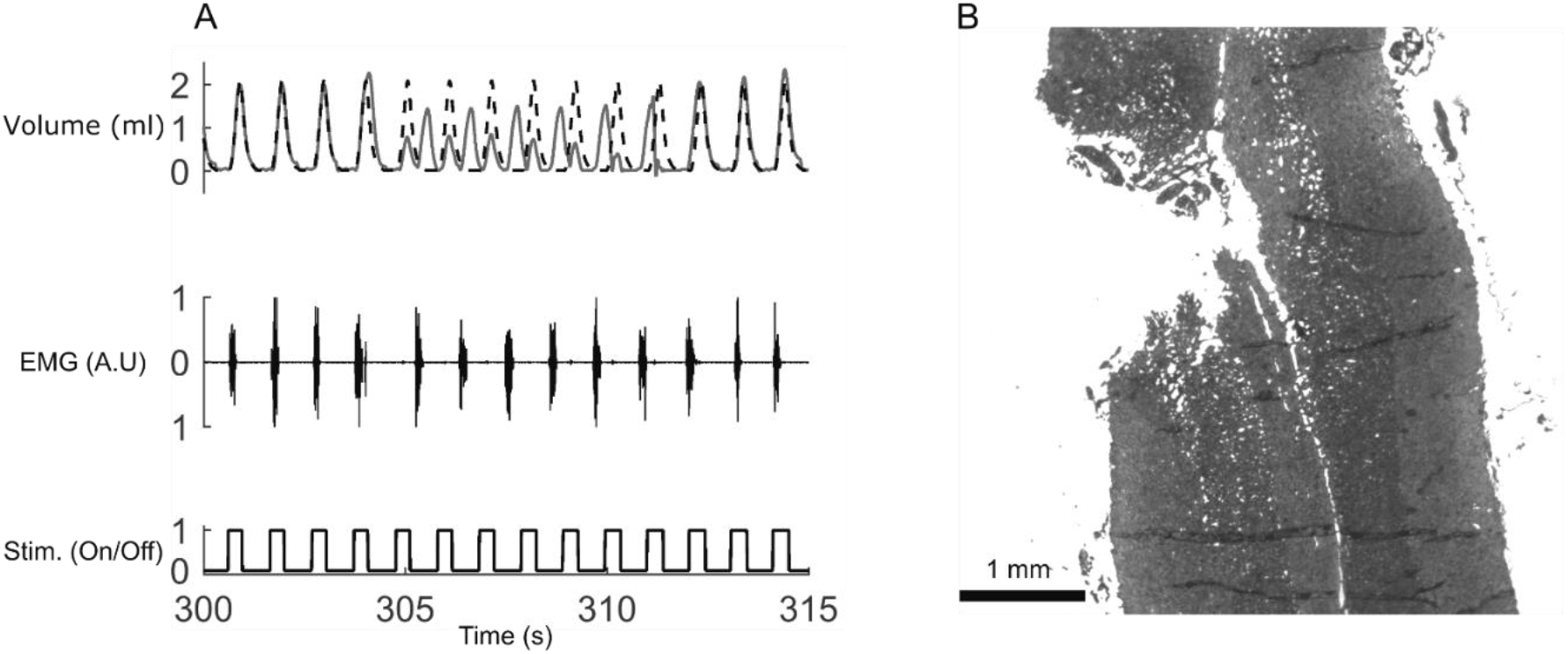
Resynchronization between intrinsic respiration and diaphragm pacing after a C2 hemisection. (**A**) Loss and recovery of synchronization between intrinsic respiration (breath volume, genioglossus EMG) and diaphragm pacing (stimulation on/off, bottom, stimulation onset denoted by grey dashed lines). De-synchronization leads to elevated error between measured (solid) and desired (dashed) breath volume. The controller is capable of re-establishing synchronization after de-synchronization due to phase shift timing between inspiratory timing of GG and pacing. (**B**) Sample histological section of the ventral spinal cord following C2 lateral hemisection. Spinal cord tissue of injured animals was longitudinally cryosectioned in 15μm sections and analyzed to assess exactness of hemisection. Hemisection was also assessed functionally by observing changes in tidal volume post-injury.

Synchronization between the desired and actual volume profiles is lost after the fourth pacing cycle, with the GG muscle activating half a pacing cycle after pacing occurs. However, this mismatch rapidly corrects itself after a few pacing cycles. De-synchronization causes an increase in the error signal to the controller and thus current amplitude delivered (not shown) increases.

Histological analysis confirmed injury to the descending respiratory pathways in all hemisected animals; a sample ventral spinal cord section from one animal is shown in Figure 9(B). Ventilatory function was also assessed by functional comparison of tidal volume prior to and after the hemisection. Hemisection led to an average decrease in tidal volume of 0.64 ml (p<.0001).

### Number of past activations, number of active neurons, and learning rate determine controller stability, performance, and adaptation speed

A total of 500 computer simulation trials were completed, each with a different permutation of learning rate, number of neurons, and past activations, to determine suitable combinations of these controller parameters that would allow for fast, accurate, and stable controller performance. Of these trials, a total of 39 parameter sets matched the selection criteria of mean iRMSE of less than 5% within 20 pacing cycles with a standard deviation of less than 2.5% iRMSE after 5% iRMSE had been achieved. Figure 10(A) shows these parameters along with number of pacing cycles required to lower iRMSE to less than 5%.

**Figure 10.**
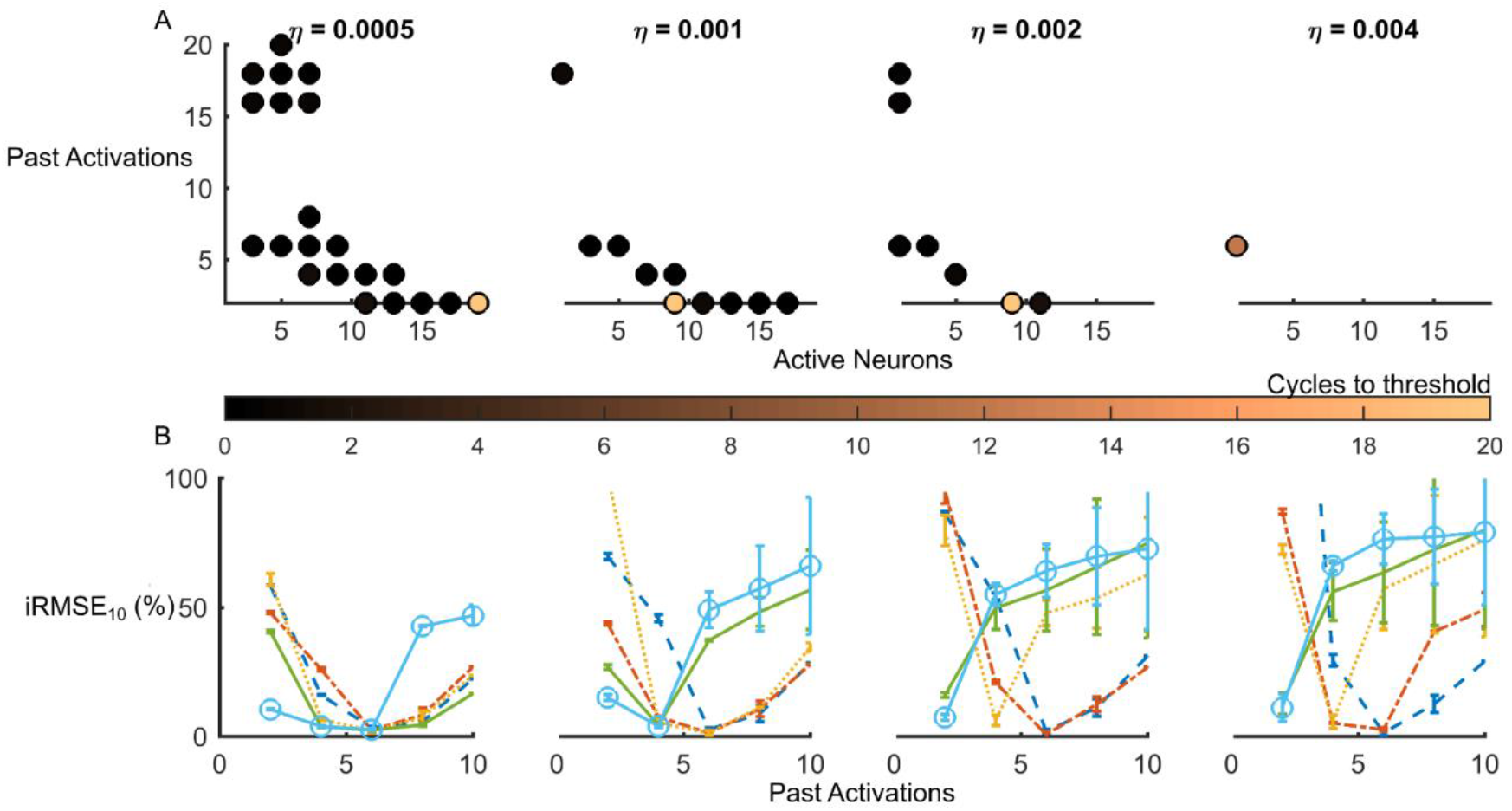
Effects of varying learning rate, number of active neurons, and number of past activations on PS controller performance as determined by simulations. (**A**) Solid dots indicate suitable parameter configurations led to adaptation that met the acceptance criteria of achieving and maintaining a 5% inspiratory RMSE within 20 pacing cycles while maintaining a standard deviation below 2.5% throughout 100 pacing cycles for several learning rates (η). Dot color denotes number of pacing cycles required to meet an inspiratory RMSE value of 5%. (**B**) Parameter trends obtained from simulation studies. Plots show effect on RMSE_10_ with respect to number of past activation values (n_p_) for several widths of the activation function given by n_a_ (1 n_a_ = blue dashed; 3 n_a_ = orange dashed dotted; 5 n_a_ = yellow dotted; 7 na = green solid; 9 n_a_ =light blue)

Results from the simulations showed that faster learning rates increased adaptation speed and thus reduce number of pacing cycles to reach the threshold. However, faster learning rates also showed a decrease in stability. As learning rate was increased, the number of acceptable parameter combinations decreased from 21 sets to only one set. The iRMSE standard deviation values of the last ten paced cycles of each trial (iRMSE10), which are assumed to be inversely proportional to stability, also showed an overall increase as learning rate increased (Figure 10(B)). When selecting a learning rate to use in the experiments, it is necessary to consider that decreased stability could pose a problem given the inherent variability of the system and presence of non-ventilatory behavior. Thus, for *in-vivo* verification of the controller parameters, a learning rate that is low enough to provide stability, yet high enough to provide sufficient adaptability, is desired.

These simulations determined a suitable range of parameters for various learning rates. For the *in vivo* studies, a low learning rate capable of maintaining robustness and stability is desired to prevent errors from propagating. Thus, a learning rate of 0.001 was deemed suitable. For this learning rate, 3 to 5 active neurons and 4 to 8 past activation values, were considered suitable parameter values as they had the lowest values for iRMSE10 % while still showing low standard deviation values. These ranges were found under the assumptions that muscle mechanics remain constant over time, i.e. that there was no fatigue, and that the breath volume profile selected as the desired trajectory is an ideal trajectory.

The controller parameters that were ultimately implemented in the intact animal studies differed from those found computationally as the presence of an intrinsic respiratory drive affected controller performance. This required controller parameter tuning to improve synchronization and entrainment with the intrinsic respiratory drive. However, the controller parameter trends that were observed computationally were generally maintained across animals. Controller parameters used can be found in Appendix Table 2

## Discussion

The present study presents a novel neuromorphic closed-loop, adaptive controller for ventilatory pacing capable of modulating stimulation parameters to elicit a breath volume profile that follows a desired profile over hundreds of breaths. A computational platform facilitated development of the controller and provided a range of controller parameters that achieved satisfactory performance, accuracy, and stability. Evaluation in intact and cervical hemisected rats validated the parameters determined computationally. The Pattern Shaper (PS) controller successfully reduced the error between measured and desired breath volume profiles in the presence of a competing intrinsic respiratory drive and/or diaphragm muscle fatigue. In hemisected rats, the PS controller successfully restored breath volume to levels observed prior to the injury while also reducing etCO_2_ values. These studies support the use of the closed-loop adaptive PS controller for ventilatory pacing to achieve breath-by-breath control after spinal cord injury or central hypoventilation syndrome.

### Computational studies provided a testbed for controller development and characterization

The simulations carried out using biomechanical computational models, described in the supplementary material, illustrate the benefits of utilizing computational models as test-bed platforms for the development of neurotechnological algorithms prior to utilization in *in vivo* studies. Here, the computational models ensured that the neuromorphic algorithm for ventilatory control was able to properly adapt and maintain a desired breath volume profile before any animal study was performed, substantially decreasing development time and use of experimental resources. These computational studies identified ranges of controller parameter values that resulted in fast, accurate, and stable performance and identified interactions between these parameters that affect adaptation performance.

The PS controller determines the amplitude of upcoming stimulation pulses as the weighted summation of the output of all neurons in the network, however, only neurons that have a non-zero output at any particular instant contribute to a stimulation pulse. Since each neuron has non-zero output for a portion of the breath cycle and the output profiles are shifted in time, the width of the output profile is directly related to the number of active neurons. Low values for this parameter localizes the influence of each neuron but might also lead to unstable behavior. However, stability can be improved by selecting an appropriate value for the number of past-activations included in the summation term of the adaptation algorithm. As shown in Figure 10, when using a low value for the number of active neurons, selecting a high value for the number of past activations was more likely to meet performance specifications.

The learning rate, *η*, directly scales the change in neuronal weight in response to each error measurement. While fast adaptation is generally desired, high learning rates can produce large and rapid changes in weights that can compromise stability or produce an under-damped response. With a low value for learning rate, the controller reaches the target iRMSE more slowly, but then the system performance remains stable.

These simulations suggest that for clinical deployment the controller would benefit from a learning rate that is fast enough to adapt to provide sufficient ventilation while being slow enough to remain stable.

### Controller provided autonomous and individualized ventilatory pacing

Animals in this study varied in weight, stimulation twitch threshold, presence of injury, and ventilatory drive as defined by etCO_2_ levels. Irrespective of these differences the PS controller autonomously adapted to achieve the desired ventilatory profile for each individual. In rodents with partial diaphragm function due to C2 hemisection, the controller adapted stimulation values to elicit tidal volume profiles that matched those observed pre-hemisection. Furthermore, the controller was able to elicit the same desired breath volume profile despite differences in the un-paced breath volume. This shows that the adaptive controller can automatically personalize treatment regardless of the efficacy of the intrinsic respiratory drive after injury.

Previous animal studies (Nochomovitz *et al*., 1983; Walter *et al*., 2011; Kowalski *et al*., 2013) have shown the feasibility of ventilatory pacing, while a number of other studies have assessed viability and safety of ventilatory pacing in humans for hypoventilation after spinal cord injury (Chervin & Guilleminault, 1994; DiMarco *et al*., 2002; Zimmer *et al*., 2007; DiMarco, 2009; Onders *et al*., 2011; Posluszny *et al*., 2014; Garara *et al*., 2016). However, these animal and human studies considered only the use of pulse trains with fixed stimulation parameters. While open-loop controllers might elicit a desired tidal volume at the time of pacing calibration, this fixed stimulation waveform must be manually selected for each individual. The selected stimulation may not be suitable after muscles fatigue, possibly leading to insufficient ventilation, discomfort, or pain. The PS controller addresses this issue with its adaptive shape-defining abilities. The PS controller achieves personalized pacing by using real-time measurements of performance (volume) to adapt pulse amplitude within the pulse train, thus allowing it to activate the diaphragm to elicit the breath volume profile desired.

Additionally, in open-loop paradigms, manual adjustment of stimulation parameters is required to maintain a constant tidal volume during short and long-term changes to the muscle or electrode interface, such as muscle fatigue or electrode encapsulation (Grill & Mortimer, 1994; Akers *et al*., 1997). The PS controller continuously adapts to achieve a fixed ventilatory pattern regardless of any change that might occur. Fatigue was not directly quantified in this study, but the presence of a gradual increase in stimulation charge delivered and a constant low iRMSE throughout the trial suggests that the controller was able to account for stimulation-induced fatigue of the diaphragm muscle. This ability to adapt diminishes or eliminates the need for clinical intervention to adjust the stimulation parameters, reducing the cost, attendant time, and inconvenience to the patient. Such consistency in ventilation might also expand the range of activities that could be performed without increasing the risk of hypoventilation or discomfort.

For the controller to be clinically viable in the presence of any degree of intact respiratory drive, the paced and intrinsic breaths should be synchronized to achieve effective and efficient ventilation. Although the PS controller does not have an explicit mechanism to achieve synchronization, it was observed that the adaptive algorithm’s intrinsic adaptation to error can promote entrainment and synchronization. This behavior occurred both at the initiation of pacing and sporadically throughout pacing. Errors that result from the temporal mismatch between the elicited and desired volume profile cause an increase in stimulation output. When the increased stimulation in the paced cycle matches an intrinsic inspiratory phase, a comparatively stronger diaphragmatic contraction occurs, causing an overshoot of the peak volume elicited. The increased pulmonary stretch receptor feedback likely triggers the Hering-Breuer reflex through vagal afferents, which resets the phase of the breathing cycle and produces entrainment and synchronization (Muzzin *et al*., 1992; Simon *et al*., 1999).

Loss of synchrony was observed in both intact and hemisected rats; however, hemisected animals also showed a higher incidence of these events and required more pacing cycles to re-synchronize. In contrast with intact animals, all hemisected animals had difficulty achieving initial phase synchronization; with pacing cycles to 10% iRMSE lasting an average of 53 pacing cycles, compared to 17.5 pacing cycles in intact animals. One possible reason for difficulty in entrainment and synchronization is that pacing in intact animals targeted a breath volume that was 20% higher than baseline, increasing stretch receptor input to the respiratory central pattern generating neural circuitry (CPG) thus strengthening the coupling between the paced and intrinsic breaths. Meanwhile, the desired breath volume in hemisected animals was the same as their baseline breath volume leading to baseline levels of stretch receptor feedback to the CPG and thus weaker coupling than in intact animals. Additionally, In hemisected animals, the hemisection unilaterally disrupts diaphragm stretch receptor-mediated feedback to the CPG via phrenic afferents(Nair *et al*., 2017), possibly reducing the strength of the primary signal mediating entrainment of the intrinsic drive by the electrical pacing.

Previous studies have suggested technology that can be used to synchronize artificial ventilation with intrinsic respiratory drive or to replicate its function. These methods include development of a controllable stimulator with an update frequency higher than the stimulation frequency and a real-time processing controller (Castelli *et al*., 2017), implementation of a bio-inspired spiking neural network model that follows intrinsic respiratory rate (Zbrzeski *et al*., 2016), predictive algorithms using body temperature and heart rate (Kimura *et al*., 1992), and breath-triggering through the use of genioglossus muscle activity (Mercier *et al*., 2017). However, these approaches have not yet been sufficiently developed or investigated.

### Dynamic control of etCO_2_ can lead to further control of ventilation

The observed decrease in etCO_2_ was lower than expected, thus indicating that a greater change in overall ventilation may be required to achieve a more pronounced decay in etCO_2_. This could be achieved by automatic adjustments to respiratory rate. In the current implementation of the controller, the PG portion of the controller produced a fixed oscillatory pattern. A PG that responds to changes in etCO_2_ and adjusts the respiratory frequency could be developed, as a respiratory pattern-generator inspired model-based PG. An adaptive neuromorphic PGPS controller may be able to provide adequate ventilation over a wide range of metabolic demands.

Although ventilatory pacing has been under development for over 70 years, and has already been used clinically, there has been a lack of closed-loop controllers for ventilatory pacing. Open-loop control has the potential to lead to inadequate ventilation as the patient undergoes changes in their physiological state, such as changes in metabolic demand and postural load, and changes in the electrode-tissue interface, such as electrode encapsulation. The automation of initial stimulation parameter selection and consequent adaptation on a breath-by-breath basis allows for individualized closed-loop control of ventilatory pacing on a long-term time scale with adaptability to short-term changes. Furthermore, this adaptive closed-loop control strategy would allow for gradual reduction of stimulation following respiratory rehabilitation, eventually leading to automatic weaning, thereby decreasing time spent in rehabilitative therapy and potentially improving ventilatory outcome measures.

## Author Contributions

All work was developed and conducted at the Adaptive Neural Systems Laboratory at Florida International University. RS developed the computational models, carried out the simulations, conducted the animal studies, and processed all data. JJA devised the concept of the original PG/PS controller, provided critical feedback on the development of the controller and data analyses, and was key to the collaborative grant submission. BKH assisted in the development of the experimental studies, provided feedback on the computational work, assisted in the statistical analysis of the data. JG prepared the animals for surgery, aided in the animal experiments, and carried out the histology and the assessment of the spinal tissue. SC guided and performed the statistical analysis of the hemisected animal data. JC participated in discussions regarding the stimulation protocol for the animal studies. SR provided feedback for the stimulation, was key to the collaborative grant submission for the French participation. RJ was the senior author who designed the overall ventilatory control scheme and provided feedback throughout the work performed, and key to the collaborative grant submission. RS, JJA & RJ contributed to the primary manuscript writing with input and editing from other authors.

## Acknowledgments

The authors would like to thank C. Vale for the assistance in the animal experiments as well as the manufacturing of the electrodes, Dr. D. Fuller for insightful conversations regarding respiratory physiology after spinal cord injury, and Dr. E. Gonzalez-Rothi for training in the spinal hemisection surgical procedure.

## Authors’ Translational Perspective

Respiratory pacing can serve an alternative to mechanical ventilation in cases of respiratory insufficiency due to trauma such as spinal cord injury (SCI) or disorders such as congenital hypoventilation syndrome. Each patient, depending on their weight, requires different breath volumes for adequate ventilation and breath-to-breath modulation of ventilation to account for changing metabolic states. Current commercially-available respiratory pacing systems are open-loop and require manual calibration by a clinician or technician to set the stimulation levels to achieve the desired breath volume. We present a closed-loop respiratory pacing controller that can automatically determine the stimulation parameters needed to achieve a desired breath volume and adapt this stimulation during an on-going breath. Experimental studies in rodents show that the adaptive controller can achieve a desired breath volume profile for a wide range of body weights, even after diaphragmatic hemiparesis caused by spinal cord injury. This adaptive approach offers multiple clinical advantages over current systems. The approach allows for a streamlined automated set-up process that provides the required pacing regardless of patient specific ventilatory biomechanics. During ongoing use, there would be a reduced risk of inadequate ventilation as the controller would adapt and respond to physiological changes such as those on muscle fatigue or alteration in posture. Furthermore, the adaptive controller could be used as a respiratory rehabilitation tool to strengthen inspiratory muscles and allow automatic weaning from mechanical ventilation. Progressive decrease in stimulation levels would provide an objective measure to gauge rehabilitation progress.

## Appendix: Tables

**Appendix Table 1.**
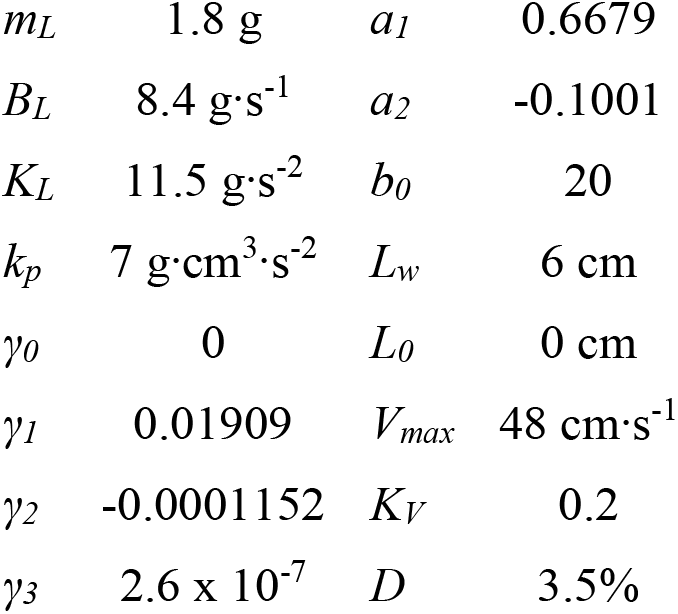
Computational model parameters

|  |  |  |  |
| --- | --- | --- | --- |
| $m_L$ | 1.8 g | $a_1$ | 0.6679 |
| $B_L$ | $8.4 \text{ g}\cdot\text{s}^{-1}$ | $a_2$ | -0.1001 |
| $K_L$ | $11.5 \text{ g}\cdot\text{s}^{-2}$ | $b_0$ | 20 |
| $k_p$ | $7 \text{ g}\cdot\text{cm}^3\cdot\text{s}^{-2}$ | $L_w$ | 6 cm |
| $\gamma_0$ | 0 | $L_0$ | 0 cm |
| $\gamma_1$ | 0.01909 | $V_{max}$ | $48 \text{ cm}\cdot\text{s}^{-1}$ |
| $\gamma_2$ | -0.0001152 | $K_V$ | 0.2 |
| $\gamma_3$ | $2.6 \times 10^{-7}$ | $D$ | 3.5% |

**Appendix Table 2.** Controller parameters that produced accurate and stable adaptaion in intact animals

| *Animal* | *Active*<br>*Neurons* | *Past*<br>*Neurons* | *Learning*<br>*Rate* |
| --- | --- | --- | --- |
| 1 | 13 | 6 | 0.0003 |
| 2 | 7 | 2 | 0.001 |
| 3 | 7 | 4 | 0.0001 |
| 4 | 7 | 4 | 0.0001 |
| 5 | 11 | 6 | 0.0003 |
| 6 | 7 | 4 | 0.001 |
| 7 | 7 | 4 | 0.001 |
| 8 | 3 | 6 | 0.001 |
| 9 | 3 | 6 | 0.001 |

## References

Abbas JJ & Chizeck HJ (1991). A Neural Network Controller For Functional Neuromuscular Stimulation Systems. In Proceedings of the Annual International Conference of the IEEE Engineering in Medicine and Biology Society Volume 13: 1991, pp. 1456–1457. IEEE.

Abbas JJ & Triolo RJ (1997). Experimental evaluation of an adaptive feedforward controller for use in functional neuromuscular stimulation systems. IEEE Trans Rehabil Eng 5, 12–22.

Akers JM, Peckham PH, Keith MW & Merritt K (1997). Tissue response to chronically stimulated implanted epimysial and intramuscular electrodes. IEEE Trans Rehabil Eng 5, 207–220.

Andrew Shanely R, Zergeroglu MA, Lennon SL, Sugiura T, Yimlamai T, Enns D, Belcastro A & Powers SK (2002). Mechanical ventilation-induced diaphragmatic atrophy is associated with oxidative injury and increased proteolytic activity. Am J Respir Crit Care Med 166, 1369–1374.

Brouillette RT & Thach BT (1980). Control of genioglossus muscle inspiratory activity. J Appl Physiol 49, 801–808.

Castelli J, Kolbl F, Siu R, N’Kaoua G, Bornat Y, Mangalore A, Hillen BK, Abbas JJ, Renaud S, Jung R & Lewis N (2017). An IC-based controllable stimulator for respiratory muscle stimulation investigations. In Proceedings of the Annual International Conference of the IEEE Engineering in Medicine and Biology Society, EMBS, pp. 1970–1973. IEEE.

Chervin RD & Guilleminault C (1994). Diaphragm pacing: Review and reassessment. Sleep 17, 176–187.

Claxton AR, Wong DT, Chung F & Fehlings MG (1998). Predictors of hospital mortality and mechanical ventilation in patients with cervical spinal cord injury. Can J Anaesth 45, 144–149.

Cnaan A, Laird N & Slasor P (1997). Using the general linear mixed model to analyse unbalanced repeated measures and longitudinal data. Stat Med 16, 2349–2380.

DiMarco AF (1999). Diaphragm pacing in patients with spinal cord injury. Top Spinal Cord Inj Rehabil 5, 6–20.

DiMarco AF (2009). Phrenic nerve stimulation in patients with spinal cord injury. Respir Physiol Neurobiol 169, 200–209.

DiMarco AF, Onders RP, Kowalski KE, Miller ME, Ferek S & Mortimer TJ (2002). Phrenic nerve pacing in a tetraplegic patient via intramuscular diaphragm electrodes. Am J Respir Crit Care Med 166, 1604–1606.

Fairchild MD, Kim SJ, Iarkov A, Abbas JJ & Jung R (2010). Repetetive hindlimb movement using intermittent adaptive neuromuscular electrical stimulation in an incomplete spinal cord injury rodent model. Exp Neurol 223, 623–633.

Faul F, Erdfelder E, Lang A-G & Buchner A (2007). G*Power: A flexible statistical power analysis program for the social, behavioral, and biomedical sciences. Behav Res Methods 39, 175–191.

Fregosi RF & Fuller DD (1997). Respiratory-related control of extrinsic tongue muscle activity. Respir Physiol 110, 295–306.

Garara B, Wood A, Marcus HJ, Tsang K, Wilson MH & Khan M (2016). Intramuscular diaphragmatic stimulation for patients with traumatic high cervical injuries and ventilator dependent respiratory failure: A systematic review of safety and effectiveness. Injury 47, 539–544.

Glenn WWL, Brouillette RT, Dentz B, Fodstad H, Hunt CE, Keens T, Marsh HM, Pande S, Piepgras DG & Vanderlinden RG (1988). Fundamental considerations in pacing of the diaphragm for chronic ventilatory insufficiency: a multi-center study. Pacing Clin Electrophysiol 11, 2121–2127.

Grill WM & Mortimer TJ (1994). Electrical properties of implant encapsulation tissue. Ann Biomed Eng 22, 23–33.

Ichihara K, Venkatasubramanian G, Abbas JJ & Jung R (2009). Neuromuscular electrical stimulation of the hindlimb muscles for movement therapy in a rodent model. J Neurosci Methods 176, 213–224.

Jackson AB & Groomes TE (1994). Incidence of respiratory complications following Spinal Cord Injury. Arch Phys Med Rehabil 75, 270–275.

John J, Fiona Bailey E & Fregosi RF (2005). Respiratory-related discharge of genioglossus muscle motor units. Am J Respir Crit Care Med 172, 1331–1337.

Jung R (2018). System and method for neuromorphic controlled adaptive pacing of respiratory muscles and nerves.; DOI: US9872989B2.

Jung R, Ichihara K, Venkatasubramanian G & Abbas JJ (2009). Chronic neuromuscular electrical stimulation of paralyzed hindlimbs in a rodent model. J Neurosci Methods 183, 241–254.

Kacmarek RM (2011). The mechanical ventilator: Past, present, and future. Respir Care 56, 1170–1180.

Kim SJ, Fairchild MD, Iarkov A, Abbas JJ & Jung R (2009). Adaptive control of movement for neuromuscular stimulation-assisted therapy in a rodent model. IEEE Trans Biomed Eng 56, 452–461.

Kimura M, Sugiura T, Fukui Y, Kimura T & Harada Y (1992). Heart rate and body temperature sensitive diaphragm pacing. Med Biol Eng Comput 30, 155–161.

Kowalski KE, Hsieh YH, Dick TE & DiMarco AF (2013). Diaphragm activation via high frequency spinal cord stimulation in a rodent model of spinal cord injury. Exp Neurol 247, 689–693.

Lessard CS (2009). Basic feedback controls in biomedicine, 1st edn. ed. Enderle JD. Morgan & Claypool.

Levine S, Nguyen T, Taylor N, Friscia ME, Budak MT, Rothenberg P, Zhu J, Sachdeva R, Sonnad S, Kaiser LR, Rubinstein NA, Powers SK & Shrager JB (2008). Rapid disuse atrophy of diaphragm fibers in mechanically ventilated humans. N Engl J Med 358, 1327–1335.

Mantilla CB, Seven YB, Zhan WZ & Sieck GC (2010). Diaphragm motor unit recruitment in rats. Respir Physiol Neurobiol 173, 101–106.

Masmoudi H, Persichini R, Cecchini J, Delemazure J, Dres M, Mayaux J, Demoule A, Assouad J & Similowski T (2017). Corrective effect of diaphragm pacing on the decrease in cardiac output induced by positive pressure mechanical ventilation in anesthetized sheep. Respir Physiol Neurobiol 236, 23–28.

Mercier LM, Gonzalez-Rothi EJ, Streeter KA, Posgai SS, Poirier AS, Fuller DD, Reier PJ & Baekey DM (2017). Intraspinal microstimulation and diaphragm activation after cervical spinal cord injury. J Neurophysiol 117, 767–776.

Milhorn HT (1966). The respiratory system. In Application of control theory to physiological systems, pp. 230–254. WB Saunders, Philadelphia, PA.

Muzzin S, Baconnier PF & Benchetrit G (1992). Entrainment of respiratory rhythm by periodic lung inflation: effect of airflow rate and duration. Am J Physiol 263, R292–300.

Nair J, Streeter KA, Turner SMF, Sunshine MD, Bolser DC, Fox EJ, Davenport PW & Fuller DD (2017). Anatomy and physiology of phrenic afferent neurons. J Neurophysiol 118, 2975–2990.

Nochomovitz ML, Dimarco AF, Mortimer TJ & Cherniack NS (1983). Diaphragm activation with intramuscular stimulation in dogs. Am Rev Respir Dis 127, 325–329.

Onders RP (2012). Functional electrical stimulation: Restoration of respiratory function, 1st edn. Elsevier B.V.

Onders RP, Elmo MJ & Ignagni AR (2007). Diaphragm pacing stimulation system for tetraplegia in individuals injured during childhood or adolescence. J Spinal Cord Med 30 Suppl 1, S25–9.

Onders RP, Ponsky TA, Elmo M, Lidsky K & Barksdale E (2011). First reported experience with intramuscular diaphragm pacing in replacing positive pressure mechanical ventilators in children. J Pediatr Surg 46, 72–76.

Peckham PH & Knutson JS (2005). Functional electrical stimulation for neuromuscular applications. Annu Rev Biomed Eng 7, 327–360.

Posluszny JA, Onders RP, Kerwin AJ, Weinstein MS, Stein DM, Knight J, Lottenberg L, Cheatham ML, Khansarinia S, Dayal S, Byers PM & Diebel L (2014). Multicenter review of diaphragm pacing in spinal cord injury: Successful not only in weaning from ventilators but also in bridging to independent respiration. J Trauma Acute Care Surg 76, 303–310.

Reynolds S, Ebner A, Meffen T, Thakkar V, Gani M, Taylor K, Clark L, Sadarangani G, Meyyappan R, Sandoval R, Rohrs E & Hoffer JA (2017). Diaphragm Activation in Ventilated Patients Using a Novel Transvenous Phrenic Nerve Pacing Catheter. Crit Care Med 45, e691–e694.

Ricard JD, Dreyfuss D & Saumon G (2003). Ventilator-induced lung injury. Eur Respir J 42, 2s–9s.

Riess J & Abbas JJ (2000). Adaptive neural network control of cyclic movements using functional neuromuscular stimulation. IEEE Trans Rehabil Eng 8, 42–52.

Sadowsky C, Volshteyn O, Schultz L & McDonald JW (2002). Spinal cord injury. Disabil Rehabil 24, 680–687.

Simon PM, Zurob AS, Wies WM, Leiter JC, Hubmayr RD, Jensen ML & Stroetz RW (1999). Entrainment of respiration in humans by periodic lung inflations: Effect of state and CO_2_. Am J Respir Crit Care Med 160, 950–960.

Stites EC & Abbas JJ (2000). Sensitivity and versatility of an adaptive system for controlling cyclic movements using functional neuromuscular stimulation. IEEE Trans Biomed Eng 47, 1287–1292.

Teixeira E, Jayaraman G, Shue G, Crago PE, Loparo K & Chizeck HJ (1991). Feedback control of nonlinear multiplicative systems using neural networks: an application to electrically stimulated muscle. In IEEE International Conference on Systems Engineering, pp. 218–220.

Walter JS, Wurster RD, Zhu Q & Laghi F (2011). Respiratory muscle pacing with chronically implanted intramuscular Permaloc electrodes: A feasibility study. J Rehabil Res Dev 48, 103.

Winslow C & Rozovsky J (2003). Effect of spinal cord injury on the respiratory system. Am J Phys Med Rehabil 82, 803–814.

Zambon M, Beccaria P, Matsuno J, Gemma M, Frati E, Colombo S, Cabrini L, Landoni G & Zangrillo A (2016). Mechanical ventilation and diaphragmatic atrophy in critically ill patients: An ultrasound study. Crit Care Med 44, 1347–1352.

Zbrzeski A, Bornat Y, Hillen BK, Siu R, Abbas JJ, Jung R & Renaud S (2016). Bio-inspired controller on an fpga applied to closed-loop diaphragmatic stimulation. Front Neurosci 10, 14.

Zimmer MB, Nantwi K & Goshgarian HG (2007). Effect of spinal cord injury on the respiratory system: basic research and current clinical treatment options. J Spinal Cord Med 30, 319–330.

